# Arousal State Control of Physiological Human Brain Pulsations

**DOI:** 10.1101/2025.01.27.635032

**Authors:** Matti Järvelä, Janne Kananen, Heta Helakari, Vesa Korhonen, Niko Huotari, Lauri Raitamaa, Johanna Tuunanen, Mika Kallio, Johanna Piispala, Hanna Ansakorpi, Vesa Kiviniemi

## Abstract

Sleep promotes cerebrospinal fluid (CSF) to interstitial fluid (ISF) exchange in brain facilitated by brain pulsations. Especially brain vasomotion and arterial pulsations modulated by noradrenaline drive the intracranial fluid dynamics. Narcolepsy type 1 (NT1) entails lessened hypocretinergic output to wake-promoting systems including the noradrenergic locus coeruleus. As arousal state and noradrenergic signaling affect CSF-ISF clearance, we chose patients with NT1 as a human hypocretin-targeted model of sleep-related pathology bridging the gap between healthy awake and sleep with respect to CSF flow pulsations. We also investigated the sensitivity of MREG to detect flow with a phantom model and sought to replicate earlier pulsation findings in sleep.

In this case-control study, we used fast fMRI to map brain pulsations in groups of healthy sleeping controls (n=13), healthy awake controls (n=79) and awake NT1 (n=21) patients. We measured the very low frequency (0.008-0.1) and cardiorespiratory frequencies and calculated in each frequency band the coefficient of variation, spectral power, and full band spectral entropy to obtain brain pulsation maps.

We uncovered a brain pulsation profile from healthy waking to sleep to a sleep-related pathology NT1 prominently affected in the vascular-related vasomotor and brain arterial pulsations. Our results established how drivers of brain hydrodynamics are affected by a specific loss of key neurotransmitter governing arousal compared to healthy sleep. We also showed with a phantom model that MREG is sensitive to flow-related signal changes and solidified evidence of brain pulsations in the healthy states of sleep and wakefulness.

**Significance statement:** This study establishes how a specific depletion of arousal state controlling neurotransmitter hypocretin-1 affects brain fluid dynamics by comparing patients with narcolepsy type 1 (NT1) to healthy wake and sleeping controls. We used fast fMRI to reveal that reduced hypocretinergic activity and following postulated inconsistent noradrenergic signaling in NT1 leads to high vasomotor and low brain arterial pulsations compared to healthy wakefulness while healthy sleep produces brain arterial and respiratory pulsations that dominate over those observed in NT1. The water flow biometrics we verified in this study with a phantom model indicate that deficient hypocretin-noradrenaline axis in humans leads to opposing changes in vasomotor and arterial induced brain pulsation that may propagate to altered glymphatic solute transportation.

## Introduction

Sleep is an integral part of human wellbeing and survival. Recent research has shown that sleep serves a housekeeping role by maintaining brain homeostasis via increased metabolic waste clearance from the brain, which may protect against various neurodegenerative disorders (1). Investigation of brain fluid dynamics has gained significant momentum after the discovery of this clearance pathway now designated as the glymphatic system. In this model, cerebrospinal fluid (CSF) enters the brain via paravascular spaces around penetrating arteries, from where arterial pulsation sieves it to the brain parenchyma facilitated by the aquaporin-4-channels (AQP4) expressed on perivascular astrocytes. Once entering the parenchyma, CSF mixes with the interstitial fluid (ISF) and exits the brain to venous paravascular spaces, along dural lymphatic vessels and the cranial nerves towards deep cervical lymph nodes (2–6). With this outflow of fluid, toxic metabolites like amyloid beta exit the brain. The precise route and the role of diffusion for fluid and metabolite clearance remains debated (7–9) but accumulating evidence from animal and human research points to an active convective fluid flow (6, 10–12).

The intracranial CSF flow is driven by the composite of three physiological pulsations: 1) intrinsic slow frequency vasomotion i.e. vasodilation/vasoconstriction and hyperemia following neuronal activation, 2) respiration-related intrathoracic pressure changes that induce rhythmic in/outflow of venous blood followed by reciprocal out/inflow of CSF, and finally 3) brain arterial pulsations induced by heart beats (13–17). Thus, it is postulated that CSF/ISF flow is facilitated by physiological brain pulsations that we have established to be measurable by fast functional magnetic resonance (fMRI) (18). In earlier fast fMRI studies, we found that the power of these pulsations increase in healthy sleep (19) and as a function of sleep depth (20).

The ascending reticular arousal system (ARAS) regulates brain arousal state via brainstem nuclei like the noradrenergic locus coeruleus (LC) (21). Tonic activation of telencephalic projections from LC neurons maintain arousal during wakefulness that switches to phasic activation during non-rapid eye movement (NREM) sleep (22). Indeed, noradrenaline release contributes memory consolidation during NREM sleep in mice, while also suppressing brain glymphatic clearance (22, 23). Narcolepsy type 1 (NT1) is a neurological disease where the lack of excitatory hypocretin-1 signaling from the hypothalamus has a direct downstream effect on the neocortex and ARAS, leading to sudden fluctuations in arousal, cataplexy and fragmented nighttime sleep (24, 25). Hypocretin-1 is a key neuromodulator for brain arousal control with high activity during normal wakefulness, a reduction during sleep and reduced/absent activity in NT1 (26). It is proposed that the absence of hypocretinergic regulation over ARAS in NT1 leads to intermitted monoamine release from brainstem nuclei like the noradrenergic LC (27, 28) making NT1 a natural human model for studying the effect of disrupted neurotransmission on the brain pulsations that drive CSF flow. In our earlier study, we had found that NT1 patients presented a mixed profile of signal variance where very low frequency (VLF) variance was increased but cardiorespiratory variance decreased compared to healthy controls (29). Yet, the relation of brain pulsations between NT1 and physiological states of healthy sleep and wakefulness remain unestablished – an inquiry that answers how neuropathological arousal state fluctuations falling in between wake and sleep caused by a loss of a specific neurotransmitter alters forces driving brain clearance.

As hypocretin-1 modulates both arousal state and noradrenergic signaling that are known to affect CSF-ISF clearance, we chose NT1 as a hypocretin-targeted model of sleep-related pathology bridging the gap between healthy awake and NREM sleep in respect of proposed CSF flow facilitated by brain pulsations. We used noninvasive whole-brain fast fMRI sequence called magnetic encephalography (MREG) to investigate brain pulsations in awake NT1 patients in contrast to healthy controls in sleeping and awake states. Furthermore, as convective flow is a prerequisite to the glymphatic theory, we use a phantom model to test the sensitivity of MREG signal to capture strictly pulsatile water flow related events. Finally, we sought to replicate our earlier results in NREM sleep, and to unify the used biometrics for brain pulsation estimation.

## Results

To compare brain pulsations between awake NT1 patients (from now on NT1 group), and healthy controls (awake HC group) during wakefulness and NREM sleep (HC NREM sleep), we employed a standardized measure of MREG time-wise signal amplitude changes (coefficient of variation: CV) and frequency domain fast Fourier transformation (FFT) spectral power calculation (spectral power: SP). We have earlier successfully applied CV to assess physiological brain pulsations in Alzheimer’s disease, epilepsy, and primary central nervous system lymphoma (30–32) but not yet in healthy NREM sleep or NT1 patients. Further, we have earlier shown that all brain pulsations exhibited increased SP in healthy NREM sleep compared to the awake state (19), but now we proceed to measure SP and spectral entropy in awake NT1 patients. Thus, we also seek to unify the biometrics used to measure brain pulsations.

### Powerful vasomotor pulsations characterize waking state narcolepsy type 1 and healthy NREM sleep

The HC NREM sleep group showed higher CV in the VLF band compared to the awake HC in a widespread bilateral spatial distribution (16 099 voxels, 435 cm^3^) covering parts of the occipital, parietal, temporal and frontal brain regions and the thalamus. We also found higher CV overlapping with parts of both lateral ventricles, superior/posterior parts of the superior sagittal sinus, and bilateral Sylvian fissure (Fig. 1A). Compared to the awake HC group, we observe that the NT1 group had higher VLF CV (3572 voxels, 96 cm^3^) in parts of the bilateral occipital cortices, the left temporal lobe, and lateral ventricle (Fig. 1A). Interestingly, we observed no significant voxels between the HC NREM sleep and NT1 groups.

**Figure 1.**
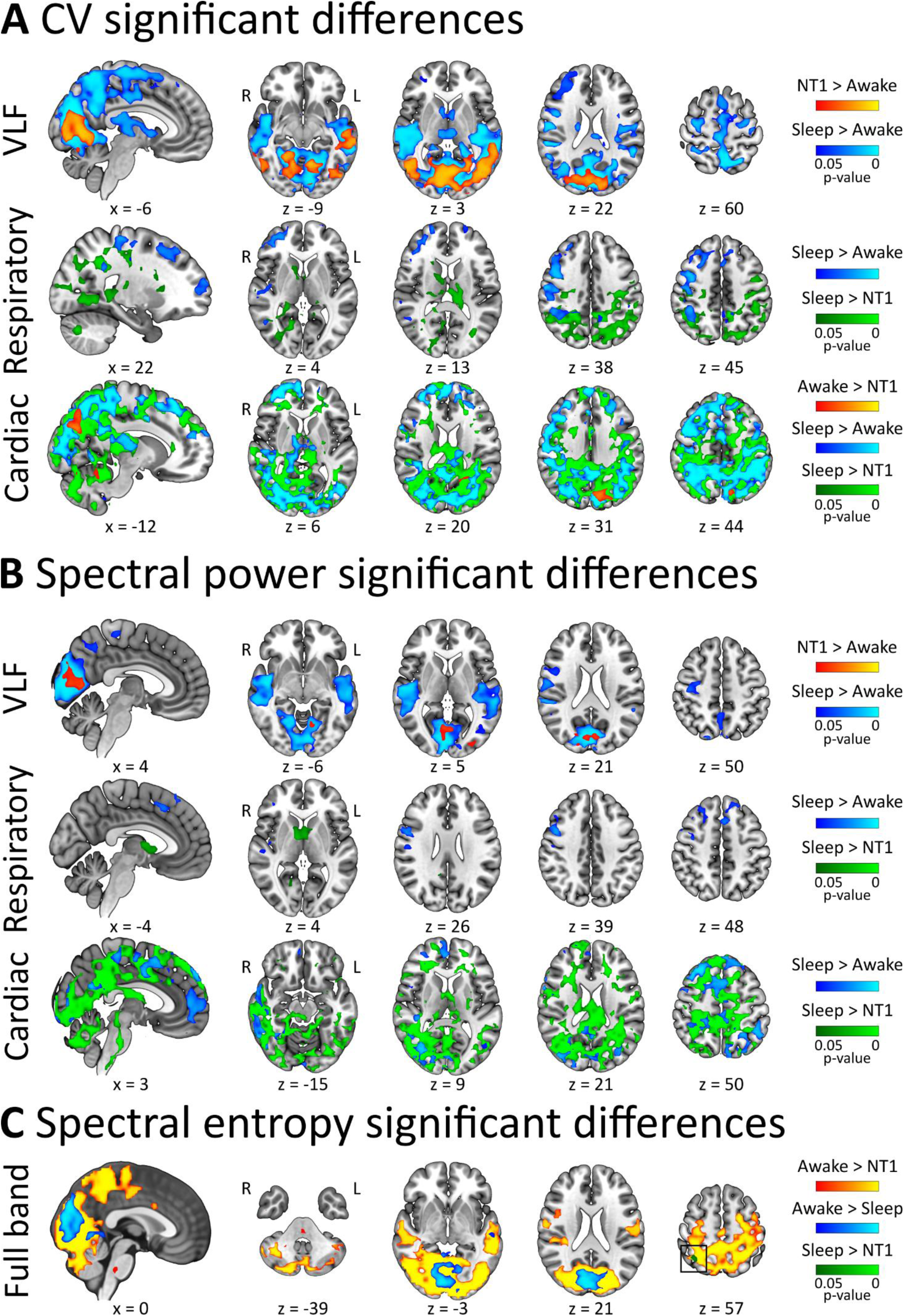
**A** CV difference maps between the three study groups in VLF and cardiorespiratory frequencies. **B** SP difference maps between the three groups in VLF and cardiorespiratory frequencies. **C** SE full band difference maps between the three groups. In the VLF pulsations, the NT1 and HC NREM sleep groups showed higher CV and SP compared to awake HC. In the cardiorespiratory pulsations, the HC NREM sleep group was characterized by highest CV and SP followed by the awake HC and finally NT1 groups. In the full band, the awake HC group showed highest SE followed by the HC NREM sleep and NT1 groups (black box highlights where NT1 SE < HC NREM sleep SE). In all comparisons, the group sizes were: NT1 n = 21, HC NREM sleep n = 13, and awake HC n = 79. VLF = very low frequency, CV = coefficient of variation, x = sagittal MNI152 coordinate, z = axial MNI152 coordinate, R = right, L = left, NT1 = narcolepsy type 1.

Compared to the awake HC group, the HC NREM sleep group had regions of higher SP in the VLF band (5239 voxels, 141 cm^3^) overlapping with parts of the bilateral occipital, sensorimotor and temporal areas of the brain and in the right somatosensory area. The higher VLF SP also overlapped with parts of the bilateral Sylvian fissure and occipital parts of the superior sagittal sinus (Fig. 1B). Compared to the awake HC group, the NT1 group showed higher VLF SP (329 voxels, 9 cm^3^) occipitally overlapping with parts of the bilateral occipital pole, cuneal, intracalcarine, supracalcarine cortices, lingual gyrus and the right lateral occipital cortex (Fig. 1B). As with VLF CV, the VLF SP did not differ between the HC NREM sleep and NT1 groups.

### NREM sleep induces the most pronounced brain arterial pulsations followed by wakefulness and lastly narcolepsy type 1

In the cardiac frequency band, we found that the HC NREM sleep group had greater CV compared to the awake HC group in widespread regions (18 000 voxels, 486 cm^3^) covering parts of the bilateral occipital, somatosensory, sensorimotor, parietal and frontal regions and right temporal areas (Fig. 1A). Further, the higher cardiac CV results encompassed parts of the frontal and occipital superior sagittal sinus, the left lateral ventricle, the right medial Sylvian fissure and inferior parts of the brainstem. Compared to the awake HC group, the NT1 group showed greater cardiac CV in a smaller occipital brain area (350 voxels, 9 cm^3^) including parts of the bilateral supracalcarine and cuneal cortices, and the left precuneous and lateral occipital cortices (Fig. 1A). Compared to the NT1 group, the HC NREM sleep group had widespread regions of higher cardiac CV (28 500 voxels, 770 cm^3^) encompassing parts of the bilateral occipital, parietal, frontal, and temporal areas. They also showed higher cardiac CV in parts of the brainstem/ARAS, around frontal and occipital parts of the superior sagittal sinus, and both lateral ventricles (Fig. 1A).

We found that cardiac frequency band SP was higher in the HC NREM sleep group compared to the awake HC group (6411 voxels, 173 cm^3^) in regions including bilateral occipital and frontal parts of the brain, and on the right side somatosensory, sensorimotor and temporal areas. This contrast also showed higher cardiac SP around the frontal parts of the superior sagittal sinus and crossing the right medial Sylvian fissure (Fig. 1B). Compared to the NREM sleep group, the NT1 group showed a large volume of lower cardiac SP (26 512 voxels, 716 cm^3^) including the bilateral occipital, sensorimotor, somatosensory, frontal, temporal and basal areas, as well as parts of the frontal and occipital superior sagittal sinus, brainstem/ARAS, the left Sylvian fissure and both lateral ventricles (Fig. 1B).

### NREM sleep in healthy controls promotes high respiratory brain pulsations, especially compared to narcolepsy type 1

In the respiratory frequency band, the HC NREM sleep group had higher CV (3042 voxels, 82 cm^3^) compared to the awake HC group in regions including parts of the bilateral frontal areas and the right somatosensory, sensorimotor and occipitotemporal areas, as well as the right medial Sylvian fissure (Fig. 1A). Compared to the NT1 group, we found that the NREM sleep group showed higher respiratory CV (5656 voxels, 153 cm^3^) in regions overlapping with parts of the bilateral occipital, somatosensory, and sensorimotor areas and the thalamus. We also found higher respiratory CV in parts of the brainstem/ARAS, and both lateral ventricles (Fig. 1A). We observed no results between the awake HC and NT1 groups.

Compared to the awake HC, the HC NREM sleep group had higher respiratory SP (854 voxels, 23 cm^3^) in parts of the bilateral frontal gyri and the right somatosensory, sensorimotor and temporal areas including part of the medial Sylvian fissure (Fig. 1B). Compared to the NT1 group, the NREM sleep group had higher respiratory SP basally, including parts of the bilateral lateral ventricles, thalamus, caudate nucleus, and the right lingual gyrus (445 voxels, 12 cm^3^) (Fig. 1B). As with respiratory CV, we found no differences in respiratory SP between the awake HC and NT1 groups.

To summarize, our findings in the time domain measured with CV together with results in the spectral domain measured with SP show that vasomotor and brain arterial pulsations differed the most between the awake HC, HC NREM sleep and NT1 groups. Notably, cardiac CV was greatest in NREM sleep, followed by healthy wakefulness, and least in the NT1 group. Vasomotor pulsations were higher in the HC NREM sleep and NT1 groups compared to the awake HC group, but interestingly did not differ from each other according to the CV and SP metrics. Respiratory pulsations were elevated in the NREM sleep group when compared to both awake HC and NT1 groups, but did not differ in the contrast between awake HC and NT1 groups. Our present findings replicate our earlier results in NREM sleep compared to wakefulness and strengthen our earlier results on signal variance differences between NT1 patients and healthy controls (19, 29).

### Spectral entropy is low in healthy NREM sleep compared to wakefulness, but even lower in narcolepsy type 1

Entropy is clinically used as an index of anesthesia depth based on the decreasing entropy as a function of anesthesia depth in electroencephalogram (EEG) (33, 34). In our prior studies, we demonstrated a decline in MREG spectral entropy (SE) as awake healthy individuals transition into NREM sleep, with an even steeper decrease associated with deeper sleep stages (19, 20). This observation led us to explore here if SE differences occurred in the present context of sleep-related disorder i.e. NT1. We assessed SE between the HC NREM sleep, NT1 and awake HC groups.

We found that compared to the awake HC group, the HC NREM sleep group presented lower SE (1441 voxels, 39 cm^3^) in regions overlapping with parts of the bilateral occipital and the left superior temporal gyrus (Fig. 1C). Compared to the awake HC group, the NT1 group showed lower SE (17401 voxels, 470 cm^3^) in widespread regions encompassing parts of the bilateral occipital, sensorimotor, somatosensory, temporal and medial areas. Moreover, the lower SE extended to posterior parts of the right lateral ventricle, lower parts of the brainstem/ARAS, and posterior parts of the superior sagittal sinus (Fig. 1C). Compared to the NREM sleep group, the NT1 group has lower SE confined to a small area (28 voxels, 0.8 cm^3^) overlapping with parts of the right superior parietal lobule and angular gyrus (Fig. 1C).

To conclude, our results showed that SE was higher in awake HC than during healthy NREM sleep, and lowest in the awake NT1 group.

### Brain arterial pulsations within an occipital brain region showing differences across all groups exhibit high accuracy in differentiation of the three states

As the cardiac pulsations follow a distinct top-down order in posterior brain regions across all three study groups, we extracted a region of interest (ROI) mask from our overlapping cardiac CV results encompassing parts of the left precuneous and cuneal cortices (Fig. 2A). To assess the fitness of pulsations in this ROI to differentiate the three groups, and to further quantify the pulsations differences, we calculated calculated the receiver operating characteristics curve (ROC) with area under the curve (AUC) for the mean CV values.

**Figure 2.**
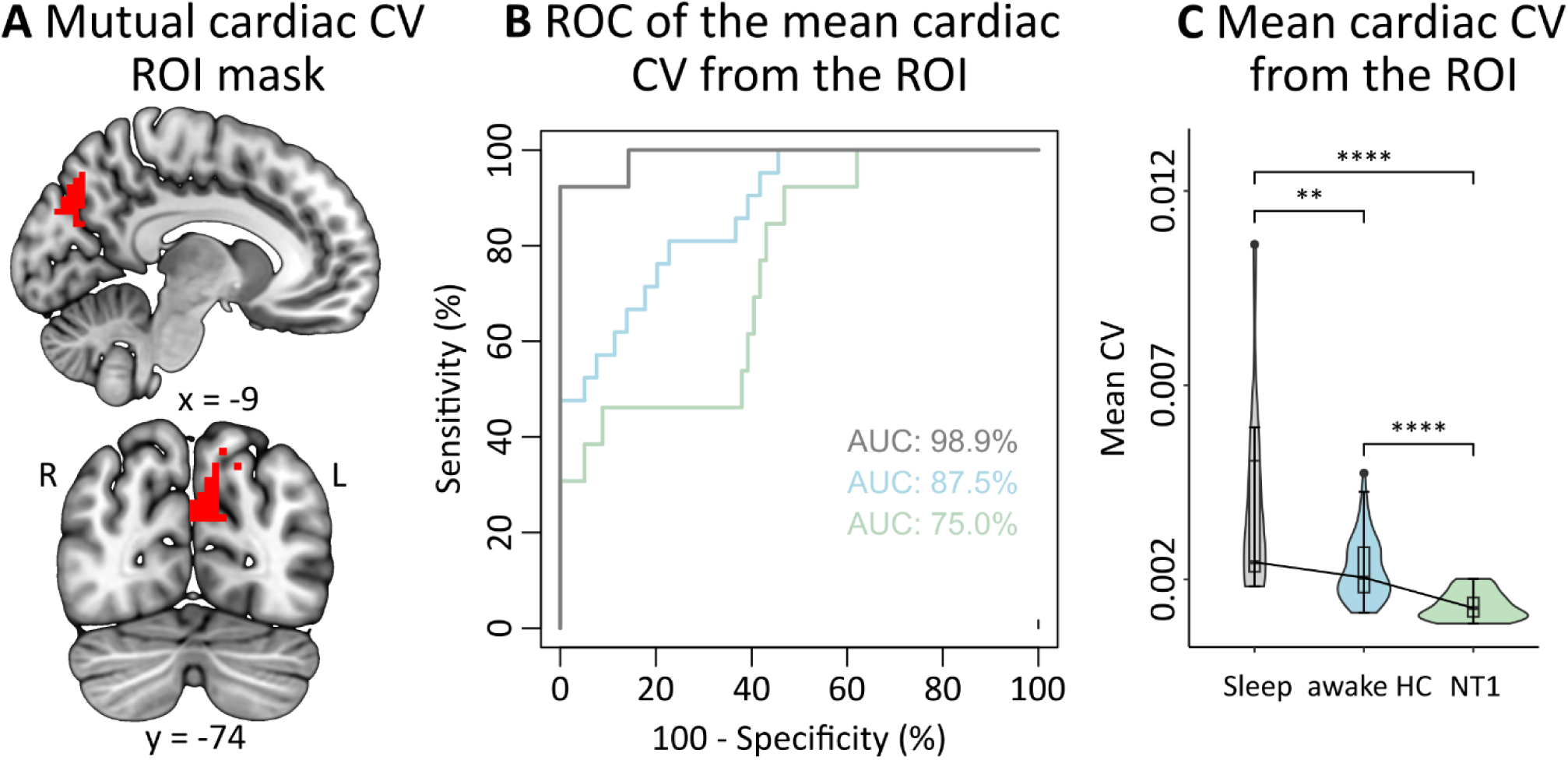
**A** The left occipital brain area common to all comparisons when comparing mean cardiac CV between the study groups (red). **B** The three groups were readily separable from mean cardiac CV values in the mutual ROI (gray: NT1 vs. HC NREM sleep, blue: NT1 vs. awake HC, green: HC NREM sleep vs. awake HC). **C** In the mutual ROI, cardiac pulsations were highest in the HC NREM sleep group, followed by the awake HC and finally by the NT1 groups. Group sizes were: NT1 n = 21, HC NREM sleep n = 13, and awake HC n = 79. HC = healthy controls, NT1 = narcolepsy type 1, CV = coefficient of variation, ROI = region of interest, ROC = receiver operating characteristics, x = sagittal MNI152 coordinate, y = coronal MNI152 coordinate, R = right, L = left.

This analysis indicated a highly accurate separation of the HC NREM sleep and NT1 groups (AUC 98.9%, Fig. 2B, grey) and good differentiation between the NT1 and the awake HC groups (AUC 87.5%, Fig. 2B, blue) as well as between the HC NREM sleep and awake HC groups (AUC 75%, Fig. 2B, green). We further found that the mean cardiac CV values in this ROI differed significantly between the three groups (Fig. 2C; Kruskal-Wallis p-value = 6.1E-09; awake HC vs. NT1 p-value = 8.8E-07; awake HC vs. NREM sleep p-value = 9.7E-03; NREM sleep vs. NT1 p = 3.4E-08).

### Pulsatile water flow induces rapid and prominent amplitude oscillations in the MREG time signal

MREG is a T2*-weighted sequence, making it sensitive if not specific to the dephasing of proton spins caused by movement of water in the static magnetic field of the scanner (35, 36). Since CSF and blood are predominantly composed of water, their flow contributes to local signal fluctuations in the T2*-weighted fMRI signals, such as MREG. To assess this sensitivity and the extent of signal fluctuations caused by water flow, we used MREG to image a phantom made from a pineapple pierced with a hole provisioned with water inlet and outlet conduits. We then employed a peristaltic pump to force water through the phantom at various flow speeds during MREG scanning. Finally, we calculated CV, SP, SE and background SP from all fluid-containing voxels at static baseline and at different flow speeds (Table 1).

**Table 1.**
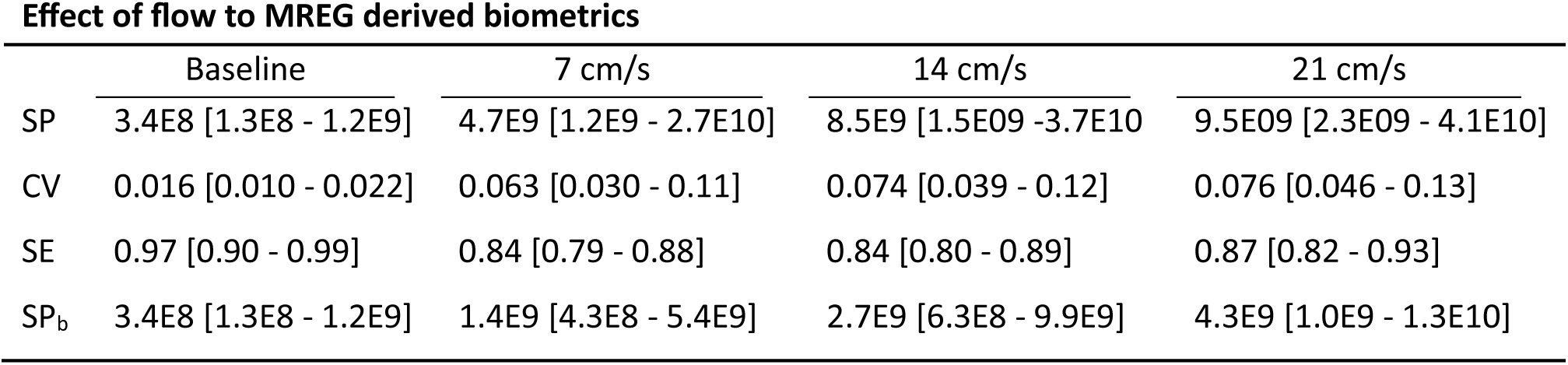
Median and interquartile range (in brackets) of SP/CV/SE/SP_b_ with different mean flow speeds. Baseline SP and SP_b_ are the same as there is no flow in the system. CV = coefficient of variation, SE = spectral entropy, SP = spectral power, SP_b_ = background SP, cm = centimeter, s = second.

Our phantom results confirmed that both SP and CV increased as a function of mean flow rate in the flow-masked ROI (Fig. 3A-B). SE showed a sharp decrease from the baseline to attain a moderately steady state between the first and second flow speeds, followed by an increase with the fastest flow (Fig. 3B). This simple phantom model demonstrates that MREG can indeed capture flow-related signal fluctuations. The spectrogram in Fig. 3A also showed higher values outside the principal 4.4 Hz peak compared to other flow speeds, hinting that an even higher sampling rate might be beneficial to capture the fastest flow events (Table 1: 12.8, 3.1 and 1.6 times greater median background SP in the 21 cm/s flow rate compared to the baseline, 7 and 14 cm/s flow rates respectively).

**Figure 3.**
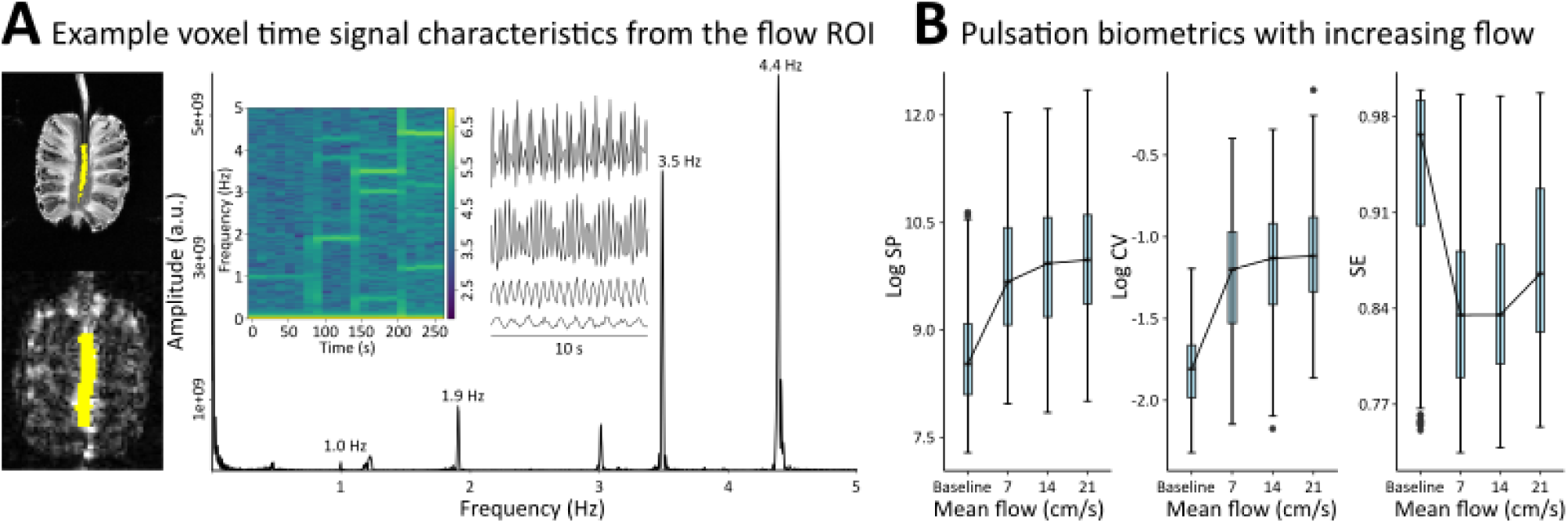
**A** Increasing water flow in the phantom model induced strong and separable T2*-weighted MREG signal amplitude oscillations (10 s signal examples, baseline as the lowest signal example and the 21 cm/s as the highest), as reflected also in the periodogram, where different flow rates correspond to distinct principal frequencies, and the 1.0 Hz peak is an known artefact from the scanner helium pump. The oscillations were also evident in the spectrogram (principal frequencies, first decree harmonics, heterodynes, and aliasing in the highest frequencies that partly exceed the critical sampling rate of 5 Hz). **B** Median biometrics measured in the frequency domain (SP) and the time domain (CV) increased as a function of water flow rate, while spectral entropy showed an initial drop and then remained stable, given that the model included only a single principal frequency event per flow rate, and finally increased due to aliasing at the highest flow rate. Semilog_10_ plots of SP and CV are presented for better visualization. A.u. = arbitrary units, cm = centimeter, s = second, Log = base 10 logarithm, Hz = hertz, SP = spectral power, CV = coefficient of variation.

**Figure 4.**
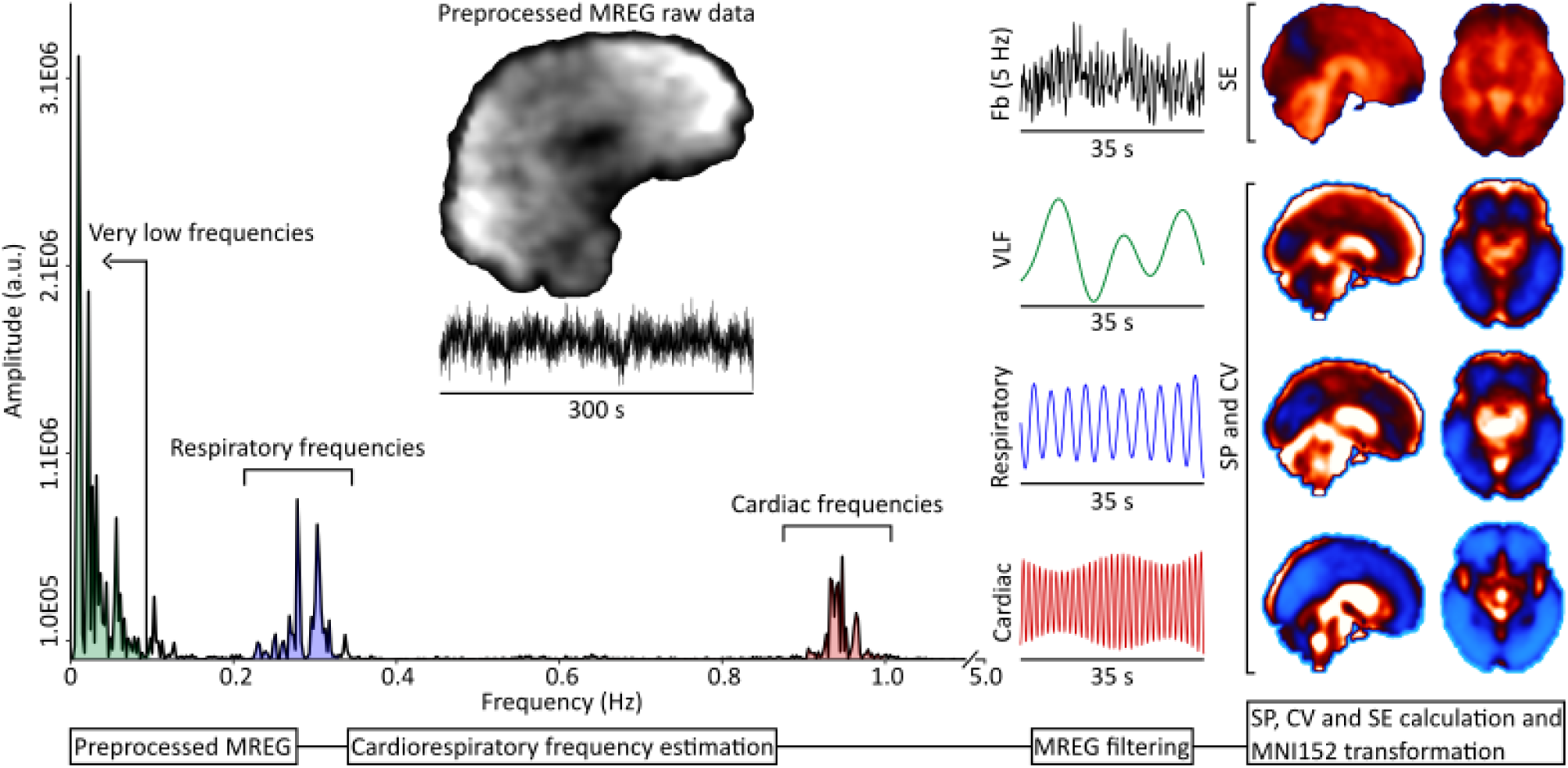
Analysis pipeline. The preprocessed MREG time domain data (black signal) was transformed into the frequency domain, where the pulsation frequencies are depicted by color (green = VLF, blue = respiratory frequencies, red = cardiac frequencies). MREG frequency domain data or peripheral physiological data was used to determine the highest peak of cardiorespiratory frequencies, and a band of 0.05 Hz on both sides of these peaks was used to filter the MREG data (i.e., to a total range of 0.1 Hz per band). Full band spectral data was used to calculate SE. VLF and extracted individual cardiorespiratory spectral data served to calculate SP maps, whereas time domain filtered VLF and individual cardiorespiratory data were used to calculate CV maps. The four brain slices on the right show that, while VLF pulsations are prominently cortical, respiratory pulsations distribute mostly to medial, ventricular and venous areas, and cardiac pulsations to ventricular and arterial areas (middle and frontal cerebral arteries areas appear bright in the axial slice). SP = spectral power, CV = coefficient of variation, SE = spectral entropy, Fb = full band, VLF = very low frequencies, Hz = hertz, s = second, a.u. = arbitrary units.

### Cardiorespiratory frequencies, blood pressure, movement and the awake HC sensitivity analysis

We did not find any group differences in the mean respiratory frequencies. Sleep is known to lower heart rate (37), and in line with this, we did find that the HC NREM sleep group has lower mean cardiac frequency compared to the awake HC and NT1 groups (Kruskal-Wallis p = 0.010, NREM sleep vs. awake HC post hoc p = 0.0074; NREM sleep vs. NT1 p = 0.040; we find no differences in awake HC vs. NT1 p = 0.72). These results further confirmed the arousal states of subjects in the HC NREM sleep and NT1 groups.

To exclude group differences in blood pressure profile that may have affected pulsation analyses (38), we compared systolic and diastolic blood pressure as well as mean arterial pressure (MAP) between the groups. We found no differences in any of these measures. To further exclude apparent pulsation findings due to head motion, we compared the mean relative and absolute motions, finding no significant group differences.

Finally, we did not find differences in CV/SP/SE maps at any pulsation frequency within the awake HC group (please see methods). This suggests that our awake HC group is rather homogenous with respect to brain pulsations, such that the differences seen in contrasts between the awake HC and HC NREM sleep groups likely reflect sleep physiology, while differences between the HC groups and NT1 group likely reflect pathology.

## Discussion

In this study, we used fast fMRI to investigate brain pulsations across the healthy awake state, NREM sleep, and in NT1 to understand how the forces driving the intracranial brain fluid flow are affected by physiological arousal states and specific pathology of arousal control. As proof of concept, we created a simple phantom model, which confirmed that the MREG sequence is sensitive to the flow of water providing a proxy of CSF and blood flow *in vivo*. We further sought to replicate in the new healthy sleeping control group our earlier findings of increased brain pulsations in NREM sleep (19), now with a significantly larger awake control group than in earlier studies.

We found that 1) NREM sleep induces high cardiorespiratory brain pulsations compared to the awake HC and NT1 groups, but interestingly there were no differences in the vasomotor pulsations between the NREM sleep and NT1 groups, 2) based on phantom measurements, water flow induces signal oscillations in the MREG data that are reflected in the pulsation biometrics, showing that MREG indeed detects flow-related events, 3) brain arterial pulsations in a posterior brain area partly overlapping with the left precuneus and cuneal cortices showed high ability to separate the three study groups, 4) our earlier findings that all brain pulsations are increased in NREM sleep compared to the awake state are replicated here, and finally 5) the same biometrics show that vasomotor pulsations are higher but brain arterial pulsations are lower in the NT1 group compared to awake HC group.

The glymphatic model proposes that CSF enters the brain parenchyma via arterial perivascular spaces and mixes with ISF, which then convey neurotoxic waste and solutes via bulk flow to and along perivenous spaces, ultimately exiting the brain to peripheral lymphatic circulation (1, 16, 17). Thus, the convective bulk flow of CSF to ISF to peripheral lymphatics is a prerequisite of the glymphatic model. The in- and outflow of intracranial CSF is thought to be driven by three distinct brain pulsations that arise from the interplay of different intra- and extracranial physiological phenomena: brain arterial pulsation waves arising from the cardiac cycle, alternating blood and CSF flow propagating from respiratory cycle, and intrinsic brain vasomotion (18).

The low frequency vasomotor waves derive from intrinsic and spontaneous reciprocal ballooning and constriction of the brain vasculature, which contribute to the regulation of local cerebral blood flow (39–41). Present findings show that NREM sleep and NT1 are characterized by high vasomotor pulsation as measured with both CV and SP, which link to CSF flow and brain clearance. Interestingly, we found no differences in vasomotor CV or SP between the NT1 and NREM sleep groups. In humans, fMRI has revealed that NREM sleep presents prominent slow-frequency oscillations in T2*-weighted blood oxygenation level-dependent (BOLD) signal, which induce an anticorrelated CSF flow in the 4th cerebral ventricle (13), which are also present in awake individuals (42). In mice, a task-induced increase in the amplitude of vasomotor frequencies correlated with tracer clearance from the brain, thus directly linking vasomotion to brain clearance (43), while sleep further increased the CSF-ISF exchange that drive tracer flux through periarterial spaces and the brain parenchyma (23). Of note, adrenergic receptor blockade in awake mice increased the tracer clearance comparable to that observed in sleep (23), thus establishing a link between brain clearance and noradrenergic signaling. Also in mice, NREM sleep manifested increased amplitude of rhythmic vasomotion of the brain pial arteries, along with reciprocal anticorrelated changes in the adjacent perivascular space (44) that is key element of the glymphatic system, which led the authors to speculate that noradrenaline may mediate this phenomenon. Indeed, in mice during NREM sleep noradrenergic signaling from the LC has been shown to drive slow vasomotor oscillations that act as a pump for brain fluid transport driving brain clearance (45). The brainstem LC neurons give rise to ascending noradrenergic projections, that comprise part of the ARAS (46). During wakefulness the LC neurons maintain high static baseline activity, which shifts to a phasic, slowly oscillating form upon sleep onset (22, 47, 48). In mice, intracerebroventricular infusion and iontophoretic application of hypocretin-1 increased the intrinsic noradrenergic cell firing in the LC (28, 49). Hypocretin-1 and -2 are excitatory neurotransmitters, which sustain cortical arousal during wakefulness through direct cortical projections arising from the hypothalamus, and indirectly via ARAS and LC (46, 50). Patients with NT1 are by definition deficient in hypocretin-1 (51). In humans, a quantitative MRI study comparing NT1 patients to healthy controls revealed lower R2 values in a brainstem area corresponding to the LC, suggesting the presence of a structural anomaly (52). Thus, the primary deficiency of hypocretin-1 in human NT1 occurs along with an LC abnormality, likely in relation to inconsistent noradrenaline release (27). We propose herein that NT1 may induce a pattern of vasomotor activity not differing from that during sleep, but distinctly higher than in the healthy awake state through secondary effects on hypocretin-1 deficiency on the regulation of noradrenergic activity, given that hypocretinergic activity is low in healthy sleep and low/absent in NT1 (26). This may imply that, even in the awake state, NT1 could promote noradrenaline-driven vasomotor oscillations akin to those seen in sleep (22). The strong wake cerebral vasomotion occurring in NT1 patients may underly their propensity to nighttime arousals and less slow wave sleep compared to healthy controls (25), but may also reflect a compensatory mechanism for the lower brain arterial pulsations in NT1 (Fig. 1A), which are normally a key driver of CSF-ISF exchange.

Brain arterial pulsations arising from the cardiac rhythm are thought to drive CSF from the periarterial spaces into the brain parenchymal ISF, thereby facilitating downstream brain fluid clearance (16). We show herein that NREM sleep leads to wide spatial distribution of high cardiac frequency CV and SP compared to the awake state, but especially compared to the NT1 group (Fig. 1A-B). Of note, the NT1 patients showed lower cardiac CV compared to awake HC, albeit on a spatially smaller scale. Given that the pharmacologically induced increase in brain arterial pulsation in mice likewise increased the CSF-ISF exchange rate facilitating brain clearance (4), present findings suggest that the interplay of CSF-ISF dynamics driven by brain arterial pulsations is most prominent in NREM sleep followed by the awake state, and lowest in the NT1 patients despite the awake state. This may relate to the hypothesized inconsistent noradrenergic release from the LC projections caused by hypocretin-1 deficiency in NT1, especially given that LC is known to regulate intracerebral vascular tone (53). In cardiac CV, we identified an occipital brain area where the three groups were readily separable (Fig. 2A-B). This ROI overlapping with part of the posterior hub of the default mode network (DMN) seems to show the greatest group differences, whereby the HC NREM sleep group presented the most pronounced cardiac CV followed with lower estimates in the awake HC and NT1 groups (Fig. 2C). In NT1, there is monotonous internetwork information propagation between the DMN and other networks (54), and the dynamic resting state activity of the DMN is unstable compared to that in healthy controls (55). Thus, this occipital ROI may represent the downstream brain area most vulnerable to the altered brain arterial pulsations.

Apart from brain vasomotion and arterial pulsations, intrathoracic pressure changes caused by the respiratory cycle promote venous blood drainage from the brain, leading to reciprocal in- and outflow of intracranial CSF (15, 56). Present results show that, compared to the awake HC, the HC NREM sleep group had higher respiratory SP, with an even wider spatial distribution of higher respiratory CV. The same holds for the comparison of the NT1 group with the HC NREM sleep group (Fig. 1A-B), but there were no differences in the contrast of the NT1 and awake HC groups. These findings confirm that, while NREM sleep is characterized by higher drive of intracranial CSF across all physiological pulsations, NT1 may impose only a minor effect on the respiratory-related flow of CSF. Our earlier study identified an increased respiratory SP in NREM sleep compared to the awake state, extending over a spatially wider area than detected in the present study. This relates to two main differences between the studies: in Helakari et. al. (19), NREM sleep and awake states were compared within the same study subjects, but here we used a larger unpaired control group and had to correct for differences in age and sex by adjusting the model. Secondly, they employed a 5° flip angle, while we now used a 25° flip angle, which increases the proportion of cardiorespiratory oscillations in the received signal (57). Despite these differences, the respiratory SP and CV results herein consistently indicate a robust increase in the respiratory pulsations in healthy NREM sleep.

The onset of sleep can be measured with entropy metrics calculated from the anesthesia monitor EEG data, where NREM sleep is characterized by low entropy (33, 34). Spectral entropy (SE) is a metric for spectral information content, which in the present context of MREG, describes how the spectral power is distributed across the sampled frequencies. The complexity of the system, here the brain, is reduced when given frequencies begin to predominate, which occurs in NREM sleep indicated by our findings both earlier (19) and in this study. We find that the NT1 group showed lower SE when compared to the awake HC group with an even wider spatial distribution than when comparing the HC NREM sleep and awake HC groups (Fig. 1C). Our SE results between the awake HC group and both the NREM sleep and NT1 groups mainly overlap with the very low frequency CV and SP results in the posterior parts of the brain, and to a lesser extent with the cardiac CV results (Fig. 1A-C). In NREM sleep and NT1, the drop in SE compared to the awake HC is then partly due to changes in the arterial pulsations, but mostly attributable to the higher vasomotor activity. This is further supported by earlier studies indicating increased low-frequency BOLD amplitude upon onset of sleep (58, 59). Thus, we suggest that the low SE that associates with high physiological brain pulsations in NREM sleep reflects a brain state characterized by optimized intracranial CSF flow to facilitate brain clearance. To our surprise, the NT1 group showed even lower SE in an area in and bordering the right angular gyrus - a posterior node of the DMN - when compared to the HC NREM sleep group (Fig. 1C). We show herein that the lack of excitatory hypocretin-1 and controlled ARAS input in NT1 lead to a local brain state with a level of spectral content usually attributed to sleep, but falling even lower than in NREM sleep.

Pulsatile flow of CSF induces signal oscillations in a T2*-weighted MRI sequence like MREG, when movement of water protons induces loss of spin phase coherence (35, 36). We find herein that increasing flow speeds are detectable in MREG time signal amplitude oscillations, power spectra and spectrograms (Fig. 3A). The increases in water flow driven by increased pulsation frequency of the peristaltic pump are accurately captured in the frequency domain by SP and in the time domain by CV (Fig. 3B). SE first decreased sharply from the resting baseline to 7 cm/s mean flow, yet remained stable from 7cm/s to 14 cm/s, while increased from 14 cm/s to 21 cm/s (Fig. 3B, Table 1). We think that the stable SE between 7cm/s to 14 cm/s is due to the singular pulsing phenomenon i.e. confining pumping power to a single narrow frequency range at a time, while the increase in SE between 14 cm/s to 21 cm/s relates to the water pumping frequency partly exceeding the critical sampling rate threshold of MREG (5 Hz), leading to both primary and aliased peaks (60). The aliasing and more turbulent flow are underlined by our findings that the median background SP in the 21 cm/s flow exceeded baseline levels by a factor of 12.8, and exceeded that at 14 cm/s by a factor of 1.6, indicating that still faster sampling may be beneficial when operating with high flow systems. We conclude that the MREG-derived pulsation biometrics calculated herein are fit to capture T2* effects related to pulsatile flow, where CV measures time signal oscillations, SP indicates spectral domain power, and SE shows the distribution of spectral content across the system.

### Strengths and limitations

The relation of physiological brain pulsations to CSF/ISF/blood flow is not without ambiguity, as real-life biological systems include many processes that may cause T2*-weighted signal oscillations. However, the T2*-weighted signal oscillations in the frequency bands that comprise cardiorespiratory brain pulsations naturally correspond to breathing and cardiac rhythm (29, 31, 61), which are known to causally induce CSF/ISF flow and blood flow in the brain (13, 16, 56). Furthermore, our phantom study confirmed that water flow has a major impact on the T2*-weighted signal oscillations. This strengthens our propositions that MREG-derived brain pulsations indeed correlate with these known physiological events.

The usual treatment for NT1 entails arousal enhancing medication, which may influence brain pulsations. As NT1 is rare and cessation of medication worsens the symptoms, we chose to include medicated patients in our study group. Because the medication regimes were heterogenous, and the NT1 group consisted of medicated and nonmedicated patients, the overall effect of medication should dilute by averaging. As such, although medication may introduce some confounding effect, the NT1 pathology common to all patients would have predominated over individual treatment effects.

NT1 is characterized by sudden shifts in the arousal state, and even falling asleep when no external input is received. In this regard, we note that: 1) we verbally checked the vigilance state of the patients at the end of scanning, 2) used the first 5 minutes of the scan, when the patients were most alert, 3) showed that ROC analysis separated the HC NREM sleep and NT1 groups with high accuracy, 4) especially the cardiac pulsations were in the opposite direction to that expected if the NT1 group patients were indeed sleeping, and 5) there were no differences in mean cardiac frequency between the NT1 patients and awake HC, but as would be expected, the HC NREM sleep group had significantly lower heart rate compared to the NT1 and awake HC groups. All these considerations suggest that the NT1 group patients were awake during scanning, although EEG is required to confirm the status.

The lessened hypocretinergic signaling in NT1 affects cortical arousal via direct effects, and via secondary impairment of the ARAS, which includes contributions from the noradrenergic LC and also ascending dopaminergic, cholinergic and histaminergic pathways (46). Noradrenaline strongly inhibits glymphatic clearance via activation of adrenergic receptors (23), and slow oscillation in noradrenaline release during NREM sleep causes reciprocal vasoconstriction and dilation, i.e., vasomotion that is anticorrelated with the diameter of the associated paravascular space (22, 44, 48). As noradrenaline strongly affects vascular tone, and given that our results in NT1 were most pronounced in the brain arterial and vasomotor frequencies, we postulate that our results mostly originate from the primary lack of hypocretin-1 leading altered noradrenergic signaling from the LC, although other monoamine systems may prove to influence brain pulsations.

The route for metabolic waste clearance from the brain remains a field of active investigation, with likely involvement of several different pathways (5–7). While the route of clearance remains debated, the present investigation of the forces driving the flow of intracranial fluids that propagate to CSF clearance is indifferent to the exact route.

### Conclusions

Present findings confirm that intracranial fluid dynamics depend heavily on the state of arousal, and engagement of the hypocretinergic system and the ARAS. We propose that the hypocretin-1 deficiency of NT1 may propagate to slow rhythmic oscillation of the brain vasculature similar to that seen in healthy NREM sleep, which persist even during wakefulness, likely driven by loss of hypocretinergic cortical excitation and by a secondary mechanism of slowly intermittent noradrenergic output from the LC (22, 44). Thus, especially vasomotor and brain arterial pulsations were affected by NT1 compared to the awake HC, but unlike NREM sleep, NT1 further leads to prominently lower brain arterial pulsations. Since both vasomotion and arterial pulsations drive the CSF-ISF exchange, we suppose that the high vasomotor activity seen in NT1 compensates for the low arterial pulsations. This phenomenon may help to explain the low cerebral amyloid burden in elderly patients with NT1 (62), insofar as high awake vasomotor activity may compensate for the suggested altered CSF clearance in NT1 (29). Finally, we verified that MREG biometrics reflect flow-related events, and replicated our earlier findings in NREM sleep compared to healthy wakefulness, which substantiates the robustness of the method for detecting the underlying physiological phenomena.

## Methods

### Participants

We conducted a retrospective registry analysis utilizing electronic patient records from Oulu University Hospital, focusing on patients diagnosed with narcolepsy, thereby identifying 66 matching cases. Among these, we successfully recruited 23 individuals diagnosed with NT1 to the study. Confirmed NT1 diagnosis with cataplexy was used as inclusion criteria. Potential confounding brain-related conditions were ruled out through screening of clinical history and by examination of structural T1 magnetic resonance images (MRI) by a neuroradiologist. One patient was excluded due to failed off-resonance correction of MREG data, and one patient was excluded due to a low mean respiratory frequency (0.11 Hz), partially overlapping with the cut-off frequency for VLF (0.1 Hz). Thus, we used data from 21 NT1 patients (NT1 group: 28.1 ± 9.2 years, 12 females) in the analyses. Among these, 15 had had CSF hypocretin-1 sampling, which ranged from 0 to 185 pg/ml. The NT1 diagnosis for the patient with 185 pg/ml was later confirmed in a specialized sleep center after progression of the disease and was thus included in the NT1 group. The remaining patients without CSF-sampling had NT1 diagnosis with reported cataplexy indicating hypocretin-1 deficiency (51). Three patients were unmedicated, and 18 had medication for daytime sleepiness and cataplexy.

For the sleeping group, we recruited 20 healthy controls from the general population of university students by email advertisement. The subjects were healthy and without self-reported continuous medication. A neuroradiologist examined the structural T1 MR images for any brain-related findings. The final sleep group consisted of 13 subjects (HC NREM sleep group: 30.5 ± 9.84 years, 6 females) with five minutes of sufficient (at least 70% of the epochs) N1/N2 sleep as confirmed by EEG (mean across subjects: 94% epochs of N1 or N2 sleep) during their MREG scanning.

We recruited 85 healthy controls with no self-reported continuous medication from the general population by advertisement. Among these, three were excluded for excess head motion (each with motion exceeding half the voxel size, please see MREG and EEG preprocessing below), and three were excluded due to low mean respiratory frequency (0.095 Hz, 0.071 Hz, 0.12 Hz) that partially overlapped with the upper VFL cut-off frequency. This led to a final group size of 79 (awake HC group: 37.3 ± 15.7 years, 52 females).

All human data was collected between 03/2018 - 08/2021. All participants gave written informed consent. This study was approved by the Ethical Committee of Medical Research in the Northern Ostrobothnia District of Finland and was conducted in accordance with the declaration of Helsinki.

### Data acquisition

All participants were scanned in Oulu University Hospital with a Siemens Magnetom Skyra 3 T MRI scanner (Siemens Healthineers, Germany), using a 32-channel head coil and fast fMRI sequence MREG. MREG is a single-shot three-dimensional (3D) sequence that employs a spherical stack of spirals undersampling the 3D k-space (63, 64). MREG data were reconstructed by L2-Tikhonov regularization with lambda = 0.1, with the latter regularization parameter determined by the L-curve method (65). An interscan crusher gradient was set to 0.1 to optimize sensitivity for physiological signals and to prevent slow signal drifts from stimulated echoes. MREG includes a dynamic off-resonance correction in k-space that corrects for respiration-induced dynamic field-map changes in the fMRI using the 3D single-shot technique (66). A T1-weighted Magnetization Prepared Rapid Acquisition with Gradient Echo (MPRAGE) scan was acquired for MREG data registration. EEG, End-tidal CO2 monitoring, photoplethysmogram, and scanner physiological data were recorded simultaneously with the MREG acquisition in a multimodal manner as described by Korhonen et al (67).

The subjects in the NT1 and awake HC groups were instructed to lie still and awake with eyes fixated on a cross in a computer screen for the whole duration of the scan. The HC NREM sleep group subjects were instructed to lie eyes closed and to fall asleep at will. Soft pads over the subjects’ ears with additional earplugs were used to minimize head motion and to block scanner noise. The arousal state of the subjects was checked after the scanning by verbal inquiry. The HC and NT1 groups were scanned in the afternoon starting at 4:00-6:00 P.M. and the HC NREM sleep group at 22:00 PM. The scans of the NT1 and HC NREM sleep group spanned 10 minutes, and the scans of the awake HC group from 5 to 10 minutes.

For the phantom model, we used a Siemens Magnetom Vida 3 T MRI scanner with 64-channel head coil, due to withdrawal from service of the Siemens Magnetom Skyra 3 T MRI machine used for human studies. A hole of 6 mm diameter was drilled through a pineapple, and tubes were connected to both sides of the hole. We then used a peristaltic pump (Watson Marlow 313S) to fill the tubes and the hole with water. We made MREG recordings during four states: 1) baseline with inactive pump, 2) mean pulsatile water flow of 7 cm/s, 3) mean flow of 14 cm/s and 4) mean flow of 21 cm/s. We applied identical imaging parameters as with the human data.

EEG of the NREM sleep group was recorded with a 256-channel high density net, using the Electrical Geodesics MR-compatible GES 400 Magstim system. Electrode impedances were confirmed to be <50 kΩ, the sampling rate was 1 kHz, and high-pass filtering was set to 0.01 Hz. Signal quality was tested outside the scanning room by recording 30 s epochs of data with eyes open and eyes closed. All EEG channels were visually checked.

### MREG and EEG preprocessing

The Oxford Centre for Functional MRI of the brain (FMRIB) software library (FSL) (68) FEAT pipeline was used for MREG data preprocessing. The data were high-pass filtered with a cut-off frequency of 0.008 Hz. The first 180 time points were excluded to minimize T1-relaxation effects. To optimize the amount of sleep data, the five minutes of EEG-recordings containing the most sleep was chosen for each HC NREM sleep group subject, and the corresponding MREG data (2861 full brain samples per subject) were extracted for further analyses. For the NT1 group, the first five minutes of the entire 10-minute scans were used to ensure better vigilance. Five minutes of MREG data were used for the awake control group, regardless of the full scan time (from 5 to 10 minutes) to match that of the other groups. Motion was corrected in four layers: 1) the raw MREG amplitude data was despiked with Analysis of FunctionalNeuroimages’ (69) (AFNI) tools, 2) FSL MCFLIRT was used to correct for bulk head motion, 3) each subjects’ MCLIRT motion correction data were used to exclude subjects with any absolute motion exceeding 1.5 mm (half the voxel size), relative motion over 0.5 mm, as well as the earlier implemented thresholds described in Järvelä et al. (29) for mean absolute motion (no subject > 0.6 mm) and mean relative motion (no subject > 0.07 mm), and finally 4) the mean absolute and relative movement were compared between the groups. We used FSL Brain Extraction Tool with neck and bias field correction for brain extraction from the 3D MPRAGE images. MREG data were spatially smoothed with a 5 mm full width and half maximum (FWHM) Gaussian kernel. For later image registration purposes, we obtained FEAT’s MREG to 3D anatomical (full-search, 12 degrees of freedom) and MREG to the Montreal Neurological Institute 152 (MNI152) 4 mm standard space transformation matrices.

For the phantom model MREG data, preprocessing steps were as above, except for omission of brain extraction and despiking, as the model was stationary.

After recording, the HC NREM sleep group’s EEG data were converted to a file format suitable for preprocessing with the Brain Electrical Source Analysis Research (BESA) software (Version 7.0). Data were then preprocessed using the Brain Vision Analyzer (Version 2.1; Brain Products). Standard preprocessing cleanup strategies were used to remove MRI gradient artifacts and ballistocardiographic (BCG) artifacts. First, we segmented the data to the correct length using the trigger marks. Gradient artifacts due to static and dynamic magnetic fields were corrected using a continuous method with no downsampling, and no filtering options (70, 71). After visual inspection of the data, we removed BCG artifacts due to blood flow effects in the scalp and brain, and from the B0 field using average artifact subtraction (70). We used semi-automatic mode to confirm that all BCG peaks were correctly chosen, and manually marked any incorrect peaks. After preprocessing of the EEG data, two trained clinical neurophysiologists, scored the sleep stages according to the American Academy of Sleep Medicine guidelines for clinical sleep studies. The final sleep stages were decided by consensus.

### Cardiorespiratory data and blood pressure

The cardiorespiratory frequencies were extracted from end-tidal CO2 and photoplethysmogram data. In the absence of these, we instead used the MREG data (for respiratory frequencies: 41 awake HC, one NREM sleep group and three NT1 subjects, and for cardiac frequencies: 39 awake HC, two NREM sleep group and six NT1 subjects). As previously described, MREG data enables accurate estimation of cardiorespiratory frequencies (29, 31, 61). We used MATLAB to obtain frequency domain spectra of the physiological data, and to estimate cardiorespiratory peak (mean) values. When using MREG data for cardiorespiratory frequency estimation, the voxel-wise preprocessed time signal data were transformed to voxel-wise FFT spectra with AFNI tools. Then, voxels in the fourth ventricle and superior sagittal sinus were used to estimate the respiratory frequencies, and voxels in the anterior/middle cerebral arteries and the lateral ventricles were used to estimate the cardiac frequencies, as these regions show the most pronounced cardiorespiratory power to visual inspection. Systolic and diastolic blood pressure were measured while seated before scanning, with loss of data for nine awake HCs and one NT1 patient.

### Voxel-wise brain pulsation parameter calculations

For each subject, the MREG full band data was filtered around their cardiac and respiratory peaks (0.05 Hz on both sides of the principal peak resulting in a total range of 0.1 Hz) and to the very low frequency (VLF: 0.008 - 0.1 Hz) range, using AFNI functions. As filtering demeans the signal, we introduced the full band signal mean back to the filtered data alike to as is done in FSL preprocessing pipeline after high pass filtering. Then, we used AFNI tools to calculate voxel-wise CV maps from the filtered data by calculating the mean and standard deviation of the data, and then dividing the standard deviation by the mean. CV provides a standardized measure of dispersion for each pulsation frequency in time domain. The transformation matrices derived from FSL FEAT were used to register the calculated CV maps to 3 mm MNI152 standard space with the FSL Linear Image Registration Tool (FLIRT). The resulting brain maps were finally masked with a 3mm MNI152 binary mask to remove extraneous voxels outside the standard brain.

We calculated SP by using FSL functions to extract the cardiorespiratory and VLF bins of the corresponding pulsation range from the MREG voxel-wise FFT spectrum. We then used AFNI tools to summate all values within these pulsation ranges, resulting in voxel-wise spectral power maps. The transformations to the 3 mm MNI152 standard brain and brain masking as above were used to produce SP maps in standard space. SP is a metric that reflects how much power accumulates into different signal frequencies in the spectral domain across the imaging experiment.

The preprocessed full band MREG data (0.008 - 5 Hz) was used to calculate in MATLAB the voxel-wise spectral entropy (SE) based on Shannon entropy, producing single value voxel-wise SE maps. SE depicts the spectral information content of the system and signal complexity, where a higher SE reflects greater signal complexity, with less prominent spectral peaks. As with the CV and SP maps, we applied the transformation to 3 mm MNI152 standard brain and masking to produce voxel-wise SE maps in standard space.

All calculated pulsation parameter maps were compared between the groups with FSL *randomise* (72) using 10 000 iterations and correcting for age and sex in the design, resulting in multiple comparisons corrected p-value maps (significant at p < 0.05). For visualization, we transformed the comparisons with significant results to 1 mm MNI152 standard space, with projection upon the MRIcroGL MNI152 standard brain using the FSLeyes spline option.

As a sensitivity analysis, we tested whether the brain pulsations differed within the awake HC group, as this could affect our further analyses. Here, we split the awake HC group into two subgroups matched for age and sex (group 1: n = 40, mean age 37.6 ± 16.2, 26 females; group 2: mean age 37.1 ± 15.5, 26 females), and compared their CV/SP/SE brain maps.

### Mutual difference mask ROC analysis

We used a posterior brain area where CV differences were present in all comparisons to create a binary mask (Fig. 2A). We then calculated ROC AUC curves for each comparison. Furthermore, mean CV values were extracted from the mask and compared between the groups (Fig. 2B-C).

### Phantom model pulsation parameter calculation

To estimate how water flow influences the brain pulsation parameters calculated from the MREG signal, we calculated voxel-wise CV, SP and SE maps of the phantom, and extracted an ROI consisting of the voxels containing water. The ROI was created by manually segmenting the water-filled space inside the pineapple from a T1 structural image. We inverted the FEAT’s transformation matrix for structural to MREG images, and used the resulting matrix to register the binary ROI mask to MREG space (579 voxels, 16 cm^3^). From this mask, we extracted both time signal and spectral data (Fig. 3A). To only include steady-state data, time signals were edited to exclude from the analysis time points coinciding with accelerating pump frequency, resulting in 462 samples per flow speed. The full band CV, SP and SE were calculated as described above in the four different flow states. To investigate non-principal, i.e., the background spectral power distribution, we removed the peak frequencies per flow speed from the phantom spectral data, and calculated SP as described above. RStudio tools were used to create the example voxel spectrum, 10 s time signal examples, spectrogram and boxplots from all voxels within the mask (Fig. 3A-B). For better visualization in the boxplots, we calculated base 10 logarithms of the CV and SP values.

The final editing of all figures was done with Inkscape version 1.3.2 and GNU Image Manipulation Program version 2.10.30. In the analyses above, we used RStudio (2023.12.1), MATLAB (R2023b), AFNI (18.0.05) and FSL (5.0.9).

### Statistics

Visual estimation and the Shapiro-Wilk test were used to examine data normality. Kruskal-Wallis test and subsequently the pairwise Dunn’s Test of Multiple Comparisons with Holm-Bonferroni multiple comparison correction was applied to test for group differences in the mutual mask mean CV values, mean absolute and relative movement values, mean cardiac frequency, and to compare systolic- and mean arterial pressure values, as these measures did not follow the normal distribution. Mean respiratory frequency and diastolic blood pressure values were tested with one-way ANOVA followed by the pairwise Tukey’s Honest Significant Difference test, as these measures were normally distributed. The CV, SP and SE brain maps were compared with FSL’s *randomise*, which uses conditional Monte Carlo random permutations implementing family-wise error-corrected threshold-free cluster enhancement correction (72) that results in multiple comparisons corrected p-value maps. In all comparisons, group sizes were: awake HC (n = 79), NT1 (n = 21), and HC NREM sleep (n = 13) except for the blood pressure comparisons, where missing data led to awake HC (n = 70), NT1 (n = 20), and HC NREM sleep (n = 13). While investigating the awake HC CV/SP/SE maps, the first group had n = 40 and the second n = 39. Significant threshold for all tests: p < 0.05.

## Acknowledgments

We wish to thank all study subjects whose attendance made this study possible. We also thank all imaging personnel who participated in data acquisition and Jussi Kantola for computational administration. We wish to acknowledge CSC - IT Center for Science, Finland, for computational resources. We thank Prof. Paul Cumming of Bern University for critical reading of the manuscript. Finally, we offer our thanks to the funders of this study: Research Council of Finland (275342, 314497, 338599, Profi 3, 314497, 335720) (VKi), Jane and Aatos Erkko Foundation (210043) (VKi), The EU Joint Programme Neurodegenerative Disease Research 2022- 120 (VKi), Finnish Cultural Foundation (MJ, JK), North Ostrobothnia Regional Fund (JK), The University of Oulu Scholarship Foundation (JK), Medical Research Center Oulu (HH, JK), The Paulo Foundation (MJ), Instrumentarium Science Foundation (MJ, JK), Finnish Medical Foundation (MJ, JK), Orion Research Foundation (MJ, JK), Suomalais-Norjalainen Lääketieteen Säätiö (MJ), Maire Taponen Foundation (JK), Finnish Brain Foundation (JK, LR) and Tauno Tönning Foundation (JK, VKo).

